# Contrasting a reference cranberry genome to a crop wild relative provides insights into adaptation, domestication, and breeding

**DOI:** 10.1101/2021.06.27.450096

**Authors:** Joseph Kawash, Kelly Colt, Nolan T. Hartwick, Bradley W. Abramson, Nicholi Vorsa, James J. Polashock, Todd P. Michael

**Affiliations:** Plant Molecular and Cellular Biology, Salk Institute of Biological Sciences, La Jolla, California 92037, USA; USDA, Agricultural Research Service, Genetic Improvement of Fruits and Vegetables Lab, 125A Lake Oswego Rd., Chatsworth, New Jersey 08019, USA; P.E. Marucci Center for Blueberry and Cranberry Research, 125A Lake Oswego Rd., Chatsworth, New Jersey 08019, USA; Department of Plant Biology and Pathology, Rutgers University, 59 Dudley Rd., New Brunswick, NJ 08901, USA

**Author notes:** **Corresponding authors:** Todd P. Michael, James Polashock.

## Abstract

Cranberry (*Vaccinium macrocarpon*) is a member of the Heath family (Ericaceae) and is a temperate low-growing woody perennial native to North America that is both economically important and has significant health benefits. While some native varieties are still grown today, breeding programs over the past 50 years have made significant contributions to improving disease resistance, fruit quality and yield. An initial genome sequence of an inbred line of the wild selection ‘Ben Lear,’ which is parent to multiple breeding programs, provided insight into the gene repertoire as well as a platform for molecular breeding. Recent breeding efforts have focused on leveraging the circumboreal *V. oxycoccos*, which forms interspecific hybrids with *V. macrocarpon*, offering to bring in novel fruit chemistry and other desirable traits. Here we present an updated, chromosome-resolved *V. macrocarpon* reference genome, and compare it to a high-quality draft genome of *V. oxycoccos*. Leveraging the chromosome resolved cranberry reference genome, we confirmed that the Ericaceae has undergone two whole genome duplications that are shared with blueberry and rhododendron. Leveraging resequencing data for ‘Ben Lear’ inbred lines, as well as several wild and elite selections, we identified common regions that are targets of improvement. These same syntenic regions in *V. oxycoccos*, were identified and represent environmental response and plant architecture genes. These data provide insight into early genomic selection in the domestication of a native North American berry crop.

## Introduction

The American or large-fruited cranberry, *Vaccinium macrocarpon*, is one of only three cultivated fruit species that are native to North America. The tart fruit is valued for its many human health benefits when consumed. For example, cranberry fruit is high in antioxidants, helps prevent urinary tract infections, has anti-atherogenic effects, and helps prevent dental caries [1–6]. The U.S. is the leading producer of cranberries with production of over 359,000 metric tons in 2019. The total value of the U.S. cranberry production in 2019 was $224.8 million dollars [7]. Canada and Chile are also major producers of cranberries with annual production in 2018 of about 195,000 and 106,000 metric tons, respectively [8] with minor production in other parts of the world. The most important products marketed are sweetened-dried-cranberries (SDCs) and juice products.

*Vaccinium macrocarpon* is a member of the Heath family (Ericaceae). Although wetland-adapted, cranberries are low-growing woody perennial vines typically growing in well-drained low pH (<5.5) sandy soils that are also low in nutrients. The roots lack root hairs and are colonized with Ericoid mycorrhizae that aid in nutrient uptake [9]. The vines produce stolons that root at various points, forming solid mats of vegetation, and cultivars are clonally propagated from cuttings. The leaves are simple and ovate with blades that measure 5-17 mm in length and 2-8 mm wide [10]. *V. macrocarpon* is diploid (2 *n* = 2 *x* = 24) and self-fertile [10]. Vertical shoots called ‘uprights’ bear the flowers and developing fruit. Cranberry blooms in early summer and each upright typically bears 5-7 white to pinkish hermaphroditic protandrous flowers (Figure S1). The flowers are 4-merous with eight anthers and an inferior ovary. The inflorescence is semi-determinate, with the flowering shoot resuming vegetative growth post-flowering. The size and number of fruit that develop per upright vary depending on the cultivar and pollination efficiency. Most modern cultivars have an average mature fruit weight of about 1.5-2.5 grams. There are four air-filled locules in the mature fruit, allowing them to float during water harvesting. Like most temperate woody perennials, cranberries go through a winter dormancy period and require 1,000-2,500 hours of chilling to resume growth and bloom in the spring [11,12].

The small-fruited cranberry, *V. oxycoccos*, is similar to *V. macrocarpon* in many ways (Figure 1). It has a similar growth habit (low-growing perennial woody vines) and it thrives in similar soils and habitats as *V. macrocarpon. V. oxycoccos*, has limited commercial production in Russia, Estonia, and Lithuania, but is mostly harvested from the wild. The ripe berries are usually red and contain similar classes of phytochemicals (e.g. anthocyanins, proanthocyanidins, flavonols, etc.) as the large-fruited cranberries. Fruit size is variable, but smaller (0.6 – 1.2 cm) than *V. macrocarpon*. The native range of *V. oxycoccos* is circumboreal, including northern Europe, northern Asia and northern North America (Figure S1). One of the key differences is in ploidy level. As noted above, *V. macrocarpon* is diploid, while *V. oxycoccos* occurs as diploid (2*n* = 2 *x* = 24), tetraploid (2*n* = 48) and hexaploid (2*n* = 72) levels [13]. Diploid *V. oxycoccus* occurs only above the 51st parallel except at high elevation, such as the Columbia mountain range [14]. Isozyme and recent sequence-based data suggest the North American diploid and tetraploid *V. oxycoccos* are likely different species [15,16]. How the hexaploids fit into the overall taxonomy is still under debate. The leaves of *V. oxycoccos* are generally smaller (8– 10 mm long and 1– 2.5 mm wide) than those of large-fruited cranberry, but length varies depending on ploidy level. The leaves of the diploid representatives, that are the subject of this paper, are about 3– 5 mm long and 1– 2 mm wide. Flowering is determinate and this species does not produce flowering uprights. Rather, the flowers arise from the stolons and tend to be darker pink than those of *V. macrocarpon*.

**Figure 1.**
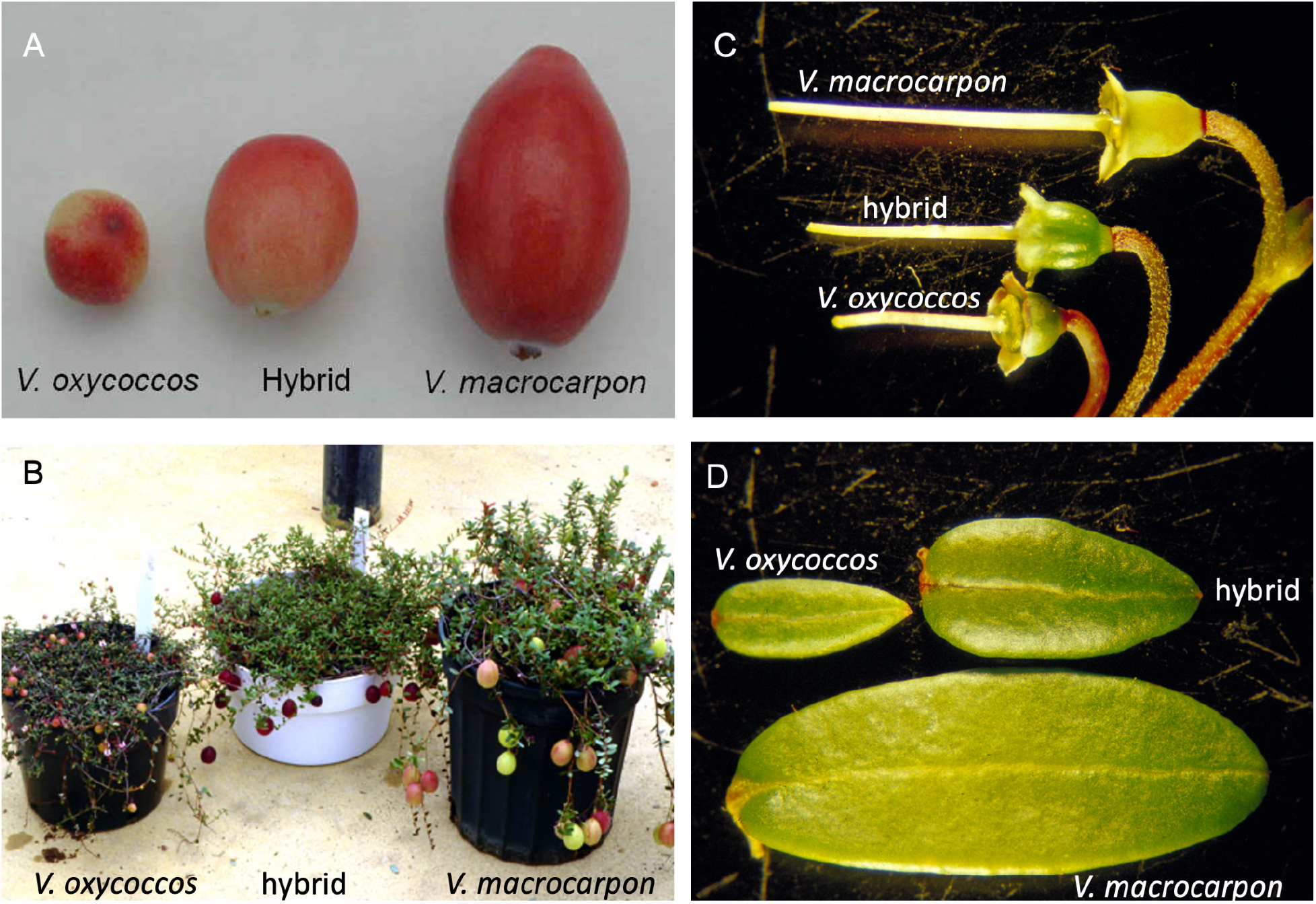
*Vaccinium macrocarpon* (Vmac) and *Vaccinium oxycoccos* (Voxy) are interfertile and have distinct morphology. A) Fruit, B) plants, C) pistils (attached to pedicels), and D) leaves from Vmac, Voxy and the interspecific F1 hybrid.

Breeding of large-fruited cranberry is in its infancy relative to most other crops, with commercial cultivars removed only one to three generations from the wild. In fact, some varieties grown today are still wild selections. The first breeding program was started in 1929 by the USDA in response to an outbreak of false blossom, a phytoplasma-incited disease. The first varieties developed in this program that were considered more ‘resistant’ to the phytoplasma (less preferable in feeding studies to the leafhopper vector of the disease) were released between 1950-1961. One of those varieties, ‘Stevens,’ is still one of the most widely grown cultivars. Breeding programs in New Jersey and Wisconsin, as well as a few private companies, are releasing new varieties with superior attributes, such as increased yield and uniform fruit color. Thirteen new varieties were released between 1994 and 2017. One focus area, particularly in the New Jersey breeding program, is fruit rot resistance. Still, all cranberry breeding to date has been limited to a single species, *V. macrocarpon*.

It is likely that in the wild, cranberries went through a genetic bottleneck during the ice age, potentially limiting variation in the available germplasm [17]. Thus, interspecific hybridization offers the opportunity to expand the genepool. *V. oxycoccos* is also reported to be very high in antioxidants and bioactive compounds, some of which may be more bioavailable than those found in *V. macrocarpon* [18]. Although there is some overlap in native range, *V. oxycoccos* is adapted to more northern latitudes, e.g., north of the 51st parallel, and may be more cold tolerant than *V. macrocarpon* (Figure S1). In addition, *V. oxycoccos*, being circumboreal, responses to photoperiod are expected to differ as this species typically has a much shorter growing season and day length varies with nearly 24 hours of light during summer. Finally, *V. oxycoccos* may offer disease resistance genes that are not found in large-fruited cranberries. Crop loss due to fruit rot remains one of the biggest challenges in the sustainable production of cranberries.

We have successfully produced F1 interspecific hybrids between *V. macrocarpon* and diploid *V. oxycoccos* and have a large F2 population segregating for many morphological, horticultural and fruit chemistry traits. However, F1 hybrids exhibit lower gametophytic fertility, e.g., lower pollen staining, than the parental species. As part of the ongoing breeding and genetics program, we are interested in comparing the genomes of these two cranberry species. We previously published a draft reference genome for the *V. macrocarpon* cultivar Ben Lear (BL) that was very useful, yet still rather fragmented [19]. Recently, a genome for a second cultivar Stevens (ST) and a fragmented assembly for *V. oxycoccos* was published [20]. Here we report an updated chromosome scale assembly for the BL reference genome and an improved draft genome of diploid *V. oxycoccos*. We further compared the genomes and transcriptomes of these two species to begin documenting their similarities and differences, as we more intensively use interspecific hybridization for domesticated cranberry improvement.

## Results

### Cranberry genome size

The first *Vaccinium macrocarpon* (Vmac) genome draft provided a resource for gene discovery and marker development [19]. The initial draft was based on a fifth generation inbred of ‘Ben Lear’ (BL-S5) and sequenced using Illumina short reads, resulting in an assembled genome size of 420 Mb and a scaffold N50 length of 4,237 bp (Table 1). We endeavoured to improve the draft genome, as well as sequence the undomesticated diploid *V. oxycoccos* (Voxy) with which we can make interspecific hybrids (Figure 1). First, we estimated the genome size of the Vmac, Voxy and the F1 hybrid (Vmac X Voxy) using k-mer frequency analysis based on short read sequence [21]. The Vmac and Voxy genomes both had single k-mer frequency peaks consistent with diploid and homozygous backgrounds with genome size estimates of 487 and 585 Mb respectively (Table 1; Figure S2; Table S1). The k-mer frequency genome size estimate for Vmac is very close to the reported flow cytometry genome size estimate of 470 Mb [22]. Conversely, the F1 hybrid had a double peak, consistent with a hybrid of two genomes that have distinct nucleotide compositions, suggesting there are distinct differences between the genomes that can be exploited for applications such as breeding (Figure S2; Table S1).

**Table 1.** Cranberry genome assembly statistics.

|  | Vmac_v1<br>(Draft_2014) | Vmac | Voxy |
| --- | --- | --- | --- |
|  | CNJ99-125-1 | CNJ99-125-1 | NJ96-20 |
| Estimated genome size Kmer19 (Mb) | 487 | 487 | 585 |
| Estimated genome size flow cytometry (Mb) | 470 | 470 | NA |
| Assembled genome size (Mb) | 420 | 484 | 484 |
| Contig (#) | 231,033 | 124 | 1,692 |
| Contig N50 length (bp) | 4,214 | 15,310,187 | 1,785,328 |
| Scaffold (#) | 229,745 | 13 | NA |
| Scaffold N50 length (bp) | 4,237 | 39,611,093 | NA |
| BUSCO | 85 | 95 | 94 |
| Repeat content (%) | 39.5 | 44.8 | 43.9 |
| Predicted genes (#) | 36,364 | 48,647 | 50,621 |
| Anchor Coverage (%) | NA | 71 | 71 |
| Anchors > 1kb | NA | 30,112 | 30,090 |
| Genome Block Coverage (%) | NA | 93 | 94 |
| Genome Block Coverage Duplicated (%) | NA | 4 | 4 |
| Genes within Block (%) | NA | 67 | 69 |
| Blocks > 100kb | NA | 412 | 412 |
| NA, not available |  |  |  |

One indicator of interspecific compatibility is fertility of the offspring resulting from interspecific crosses. Pollen in *Vaccinium* spp. is shed as a tetrad; four microgametophytes result from a meiotic event held in a tetrahedron. Staining of the tetrads with lacto-phenol blue and counting those pollen grains within each tetrad that take up the stain as potentially viable can be used as an indicator of pollen viability. Pollen from ‘Stevens’ and other commercial cultivars was estimated to be 99% viable, while accession NJ96-20 of Voxy was 56% and NJ96-127 of Voxy was 80%. This shows that there is some pollen infertility in this limited representation of Voxy. The F1 hybrids varied from about 29%-53% pollen stainability. This suggests at least some interspecific meiotic recombination leads to interlocus allelic instability in the gametophtic generation in the F1 (Vmac x Voxy) hybrids.

### Updated V. macrocarpon (Vmac) genome

Long read sequencing has enabled a new wave of near-complete plant genomes [23]. We sequenced the same fifth generation inbred (BL-S5) using Oxford Nanopore Technologies (ONT) long read sequencing, and assembled the reads using a correction-free overlap, layout, consensus (OLC) strategy [24]. The resulting genome was an extremely contiguous assembly with a total length of 484 Mb in 124 contigs and a 15 Mb N50 length, representing whole chromosome arms with few repeats in the genome assembly graph consistent with the inbred nature of the line (Vmac_v1; Figure 2A;Table 1). The genome assembly was collinear with the chromosome-scale, haplotype-resolved blueberry (*Vaccinium corymbosum*) genome (Figure S3) [25]. We scaffolded the Vmac_v1 genome into chromosomes leveraging the high contiguity of the chromosomes and the synteny with haplotype1 of blueberry, and then validated the order and orientation of the Vmac scaffolds (chromosomes) reported here using high-density genetic map markers [26]. The resulting Vmac chromosome-scale assembly was contiguous with other closely related species that have chromosome-scale assemblies such as rhododendron (*Rhododendron williamsianum*) [27], persimmon (*Diospyros lotus*) [28], tea (*Camellia sinensis*) [29] and kiwi (*Actinidia chinensis*) [30] (Figure 2B).

**Figure 2.**
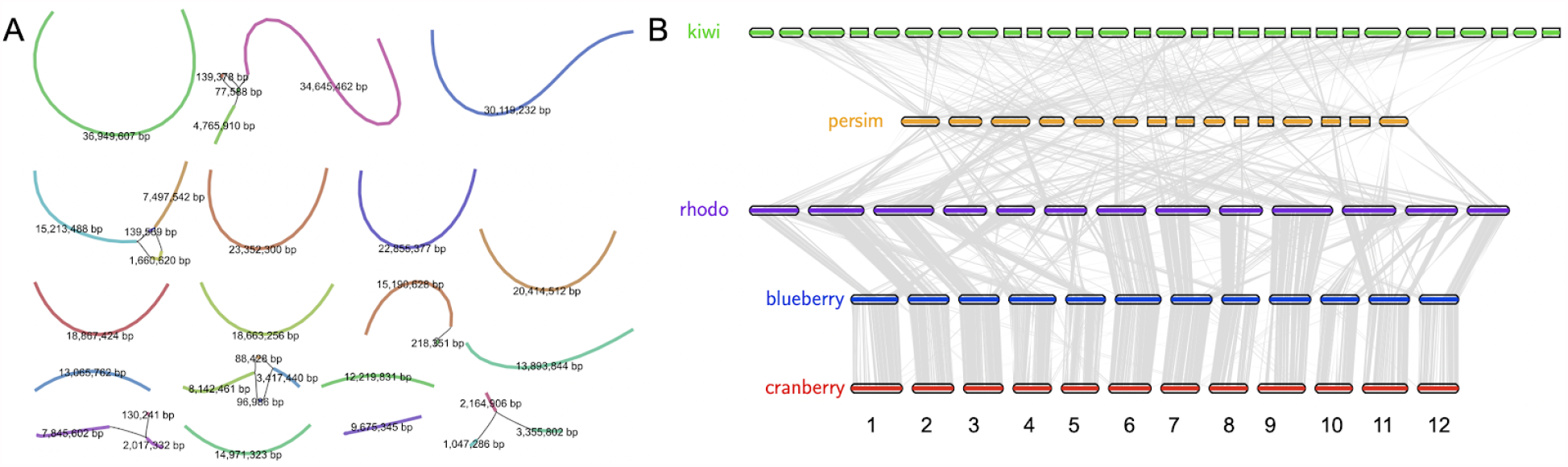
Chromosome scale assembly of *Vaccinium macrocarpon* (Vmac). A) Genome assembly graph of the cranberry genome resulted in chromosome scale, reference quality, contigs and very few “hairballs” (graph bubbles, or multiple connections due to heterozygosity and repeats). Contig size (bp) labeled, and color is randomly assigned. B) Protein based alignment of the cranberry (red; *Vaccinium macrocarpon*) genome with blueberry (blue; *Vaccinium corymbosum*), rhododendron (purple; rhodo; *Rhododendron williamsianum*), persimmon (orange; persim; *Diospyros lotus*), and kiwi (green; *Actinidia chinensis*). Lines (grey) symbolize pairwise syntenic blocks between genomes.

### The V. oxycoccos (Voxy) genome

Next, we sequenced the diploid Voxy genome using the same long read approach as with Vmac. Unlike Vmac, Voxy did not assemble into chromosome arms, yet was of high quality with a total length of 484 Mb in 1,692 contigs with an N50 length of 1.8 Mb (Table 1; Figure S4). The more fragmented nature of the Voxy assembly most likely reflects the underlying heterozygosity relative to the near-homozygous Vmac fifth-generation inbred (BL-S5). Both Vmac and Voxy assemblies were very complete with 95 and 94 BUSCO percentages, respectively (Table 1; Table S2). Repeat annotation that leveraged a *de novo* pipeline [31], predicted that 44.8 and 43.9% of the Vmac and Voxy genomes were repeat sequences, respectively (Table 1; Table S3). We generated full-length cDNA sequences using the ONT platform and used these reads to predict protein coding genes in Vmac and Voxy and found 48,647 and 50,621 respectively (Table1). We annotated the telomeres (AAACCCT), which revealed Vmac has longer telomeres (∼12 kb) than Arabidopsis (∼3 kb) [32] (Figure S5). In addition, we found centromeres with higher order repeats and base arrays of 124 bp (Figure S5).

The genomes of Vmac and Voxy are highly syntentic, with 80% of the genomes contained in syntenic blocks (Figure 3A; Table 1; Figure S6). Highly conserved syntenic gene connections are maintained between Vmac and Voxy, such as the tight linkage between the core circadian clock genes *LONG ELONGATED HYPOCOTYL 1* (*LHY*) and *PSEUDO RESPONSE REGULATOR 9* (*PRR9*) that dates back to mosses (Figure 3D). Vmac does have a tandem duplication of *LHY* that is specific to Vmac. It is not found in Voxy or blueberry (Figure S7). Within the syntenic blocks, there is 60% fractionation (4/10 genes are lost between Vmac and Voxy) and remnants of a previous whole genome duplication (WGDs) at 10% fractionation (Figure 3B). While both Vmac and Voxy maintain 5% of their genomes in 2 syntenic blocks (Figure S6), there are 54 and 70 genes that are specifically duplicated between them respectively (and retained in both). Voxy genes retained in duplicate are overrepresented for phenolic glucoside malonyltransferase that is involved with pest defense [33], providing a possible source of genes for Vmac improvement.

**Figure 3.**
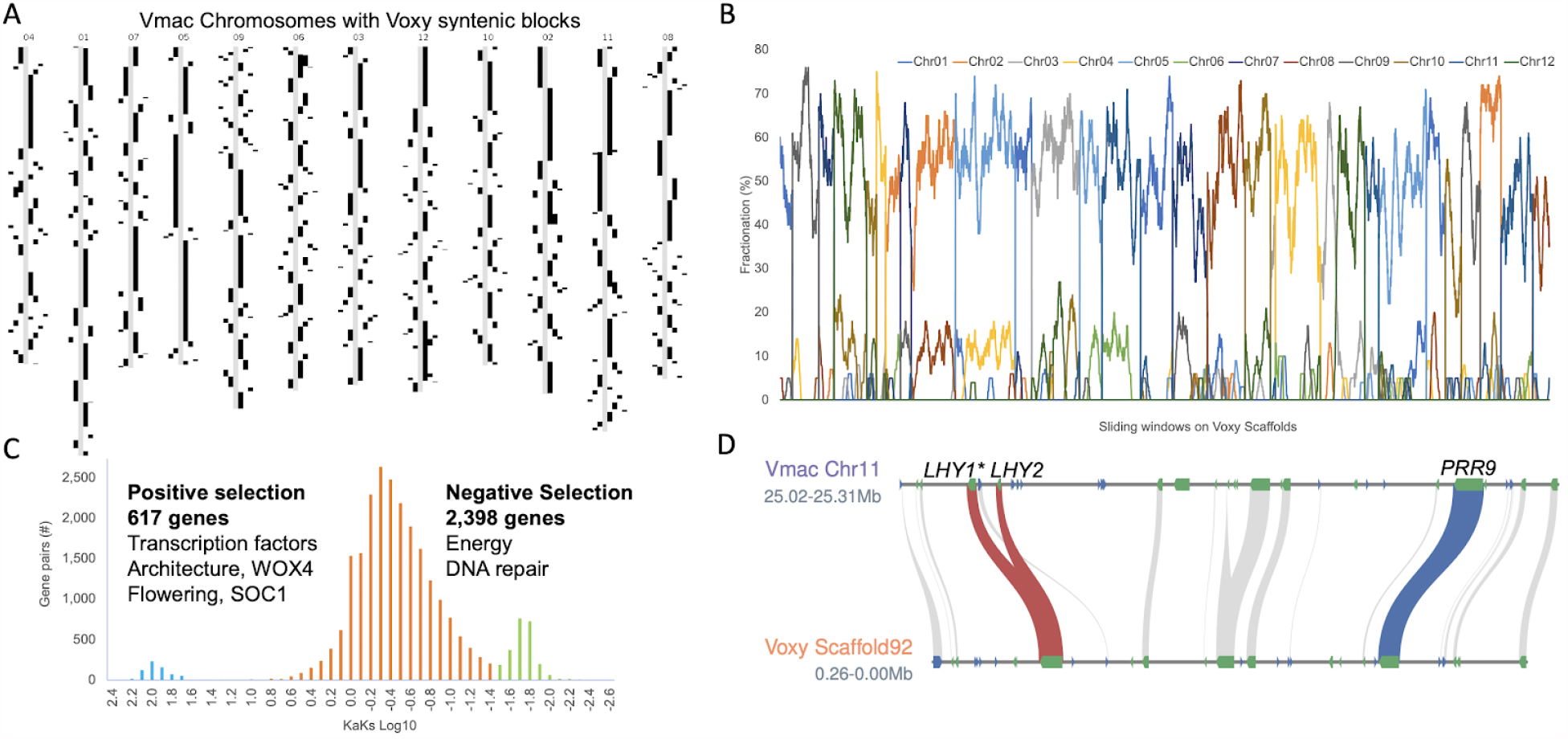
*V. macrocarpon* is highly syntenic to its wild relative *V. oxycoccos*. A) Voxy syntenic blocks (black) visualized on Vmac (grey) chromosomes. B) Fractionation of Vmac genes (chromosomes labeled in color) across Voxy scaffolds shows 2:2 syntenic depth with 60% genes present in the most recent whole genome duplication (WGD). C) Non-synonymous (Ka) by synonymous (Ks) base changes between Vmac and Voxy syntenic pairs reveals genes under positive (blue; far left) and negative (green; far right) selection. D) Conserved linkage between core circadian clock genes *LATE ELONGATED HYPOCOTYL* (*LHY*; red lines) and *PSEUDO RESPONSE REGULATOR 9* (*PRR9*; blue line) maintained on Vmac Chr11 and Voxy Scaffold92; other syntenic genes in the region (grey). Vmac has a tandem duplication of LHY (*LHY1* and *LHY2*) not found in Voxy, and under positive selection (*) in the cranberry cultivar Mullica Queen (MQ).

Vmac and Voxy have different growth habits, architecture, flower and fruit sizes, as well as distinct ecological ranges (Figure 1; Figure S1). We looked at the syntenic genes to determine those under selective pressure between Vmac and Voxy and found that there were 617 genes under positive selection (Ka/Ks>1) and 2,398 under negative selection (Ka/Ks<1) (Figure 3C; Table S4). The genes under negative selection are related to primary processes such as energy acquisition and DNA repair. In contrast, the genes under positive selection are transcription factors, architecture genes (e.g. *WUSCHEL RELATED HOMEOBOX 4, WOX4*), and flowering genes (e.g. *SUPPRESSOR OF OVEREXPRESSION OF CO 1, SOC1*) (Figure 3C; Table S4). Tandem duplications (TDs) also provide clues as to the differences between closely related species. Vmac and Voxy have 2,619 and 2,580 TD clusters, which is similar for other genomes of this size range. While many of the TDs are shared between the two species, there are 37 and 41 unique GO terms that separate Vmac from Voxy respectively (Table S5,6). Vmac specific TDs were focused on GO categories associated with plant architecture, lipid metabolism, hormone stimulus, and phenol-containing compound metabolism. In contrast, Voxy unique TDs were more focused on response to the environment (cold, wounding), toxin catabolic processes, and root development (Figure S8).

Often crop wild relatives retain disease resistance genes that are lost in a crop during domestication, resulting in the wild relatives having more or different disease resistance genes [34]. Leveraging an approach that identifies genes with disease resistance domains [35], we found that Voxy had 9,950 domains in 1,787 genes, while Vmac had 10,081 domains in 1,795 genes. 65% of the predicted disease resistance genes were shared between Vmac and Voxy in syntenic blocks, which means 35% represent presence/absence variation (PAV) between the two genomes (Table S7). Of the disease gene PAVs, 62 and 65% were in TD regions, consistent with each species having amplification of disease resistance genes specific to their genomes (Figure S9).

### Cranberry genome evolution

Cranberry differs in specific ways from its close relative, highbush blueberry (*V. corymbosum*), such as in stature (low-growing vine vs. crown-forming bush), fruit chemistry, and berry types (e.g. ripe cranberries are firm, high in proanthocyanidins, high in acidity [36], and low in sugar (< 6%), while blueberries are soft and sweet (>12%). The contrast in fruit chemistry reflects divergence in seed dispersal mechanism, i.e., abiotic (cranberries float on the water) versus animal (blueberries are eaten by birds etc. that disperse the seeds). We clustered the proteomes of Vmac, Voxy, blueberry, rhododendron, persimmon, tea, and kiwi to identify both shared and cranberry-specific genes and pathways (Figure 4A). We found 8,089 orthogroups (OGs) shared across all genomes, 6,076 specific to Vmac and Voxy, 5,223 specific to blueberry, and 3,119 shared in the *Vaccinium* spp. (Figure 4B). There are fifteen overrepresented gene ontology (GO) terms from cranberry specific orthogroups as compared to blueberry (Figure 4E; Table S8). There were also 718 and 518 OGs specific to Vmac and Voxy respectively. The overrepresented GO terms from these OGs in Voxy are pantothenate biosynthesis (vitamin B5) and plant architecture; the latter being consistent with the genes under positive selection (Table S4,8). In contrast, the overrepresented GO terms in Voxy were focused on nitrogen processes (Table S8).

**Figure 4.**
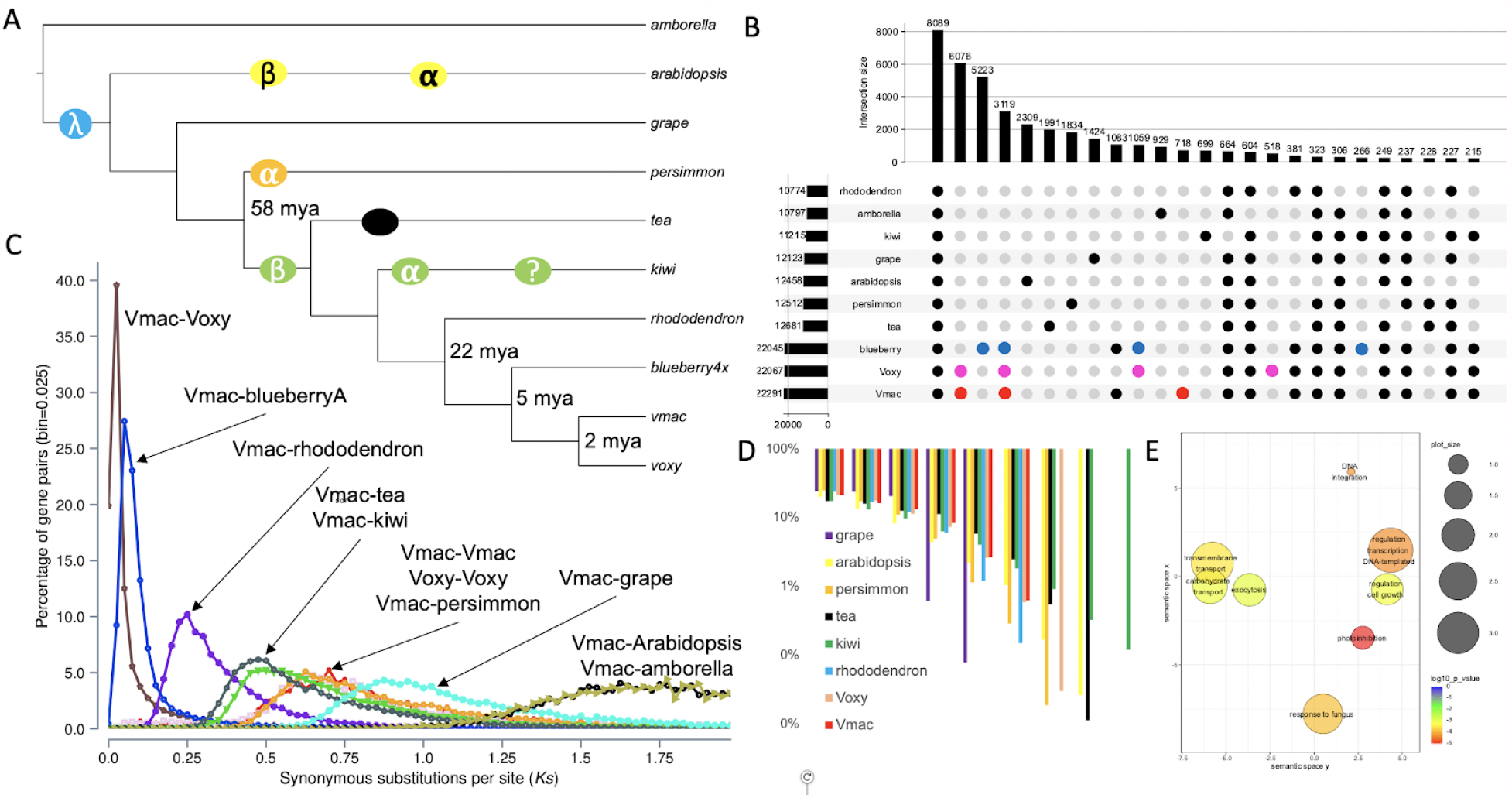
Whole genome duplication evolution of cranberry. A) Phylogenetic tree built with single copy proteins across amborella (*Amborella trichopoda*), arabidopsis (*Arabidopsis thaliana*), grape (*Vitis vinifera*), persimmon (*Diospyros lotus*), tea (*Camellia sinensis*), kiwi (*Actinidia chinensis*), rhododendron (Rhododendron *williamsianum*), blueberry 4x (tetraploid *Vaccinium corymbosum*), Vmac, and Voxy. Circles symbolize whole genome duplications (WGD) events; . B) Upset plot of the overlap between gene families. Red, pink and blue dots emphasize some of the similarities and differences among the *Vaccinium* spp. C) Synonymous substitution (Ks) distribution plot across species. D) Shared syntenic blocks compared to amborella across species. E) Significant gene ontology (GO) terms for cranberry (Vmac and Voxy) specific orthogroups (OGs) plotted in semantic space.

The chromosome-resolved Vmac genome provided the opportunity to gain a better understanding of the evolution of the cranberry genome. The recent chromosomal-scale assembly of rhododendron (*R. williamsianum*) suggested that there are two shared WGDs in the Ericales lineage [27] (Figure 4C). Consistent with Vmac containing these two WGD is the 4:1 syntenic ortholog pattern between Vmac and amborella, which is the basal plant lineage without a WGD (Figure 4D). Based on synonymous substitutions (Ks) between syntenic ortholog pairs, Vmac is separated from its two closest relatives with chromosome-resolved genomes, blueberry and rhododendron, by 5 and 22 million years ago (mya) respectively. Moreover, while Vmac and Voxy diverged 1-2 mya, the Ks for its paralogous genes suggests the most recent WGD occurred 58 mya, which is consistent with the timing of the WGD found in kiwi (Ac-***α***) and rhododendron (*Ad-**β***), but not persimmon (Figure 4A,C) that has its own WGD event (*Dd-**α***) [27,37,38]. Therefore, Vmac and Voxy contained the two WGDs *Ad-**β*** and ƛ WGDs that have shaped their genomes.

### Resequencing inbreds and parents reveal regions of selection

Cranberry is a relatively young crop in terms of years post domestication, and many high value cultivars are only modestly improved over wild selections [22]. We resequenced several cultivars important to the cranberry breeding program to identify regions of the genome that may still provide genetic resources for breeding as well as genes that are under selection in these cultivars. We looked at four cultivars that represent three generations of breeding: Stevens (ST), #35, Mullica Queen (MQ), and the Ben Lear (BL) parent. BL is a wild selection from 1901, while ST and #35 are first generation selections from crosses of wild selections, and MQ is a second generation offspring between a wild selection and #35 [26]. We also resequenced each generation from the BL-Self series (BLS1-BLS7) to identify the variation that was lost during the inbreeding process. We mapped the reads, identified SNPs between the cultivars and inbreds, and then looked for trends in variation in 250 kb bins, which highlighted the regions of the Vmac genome with high diversity (Figure 5A,B).

**Figure 5.**
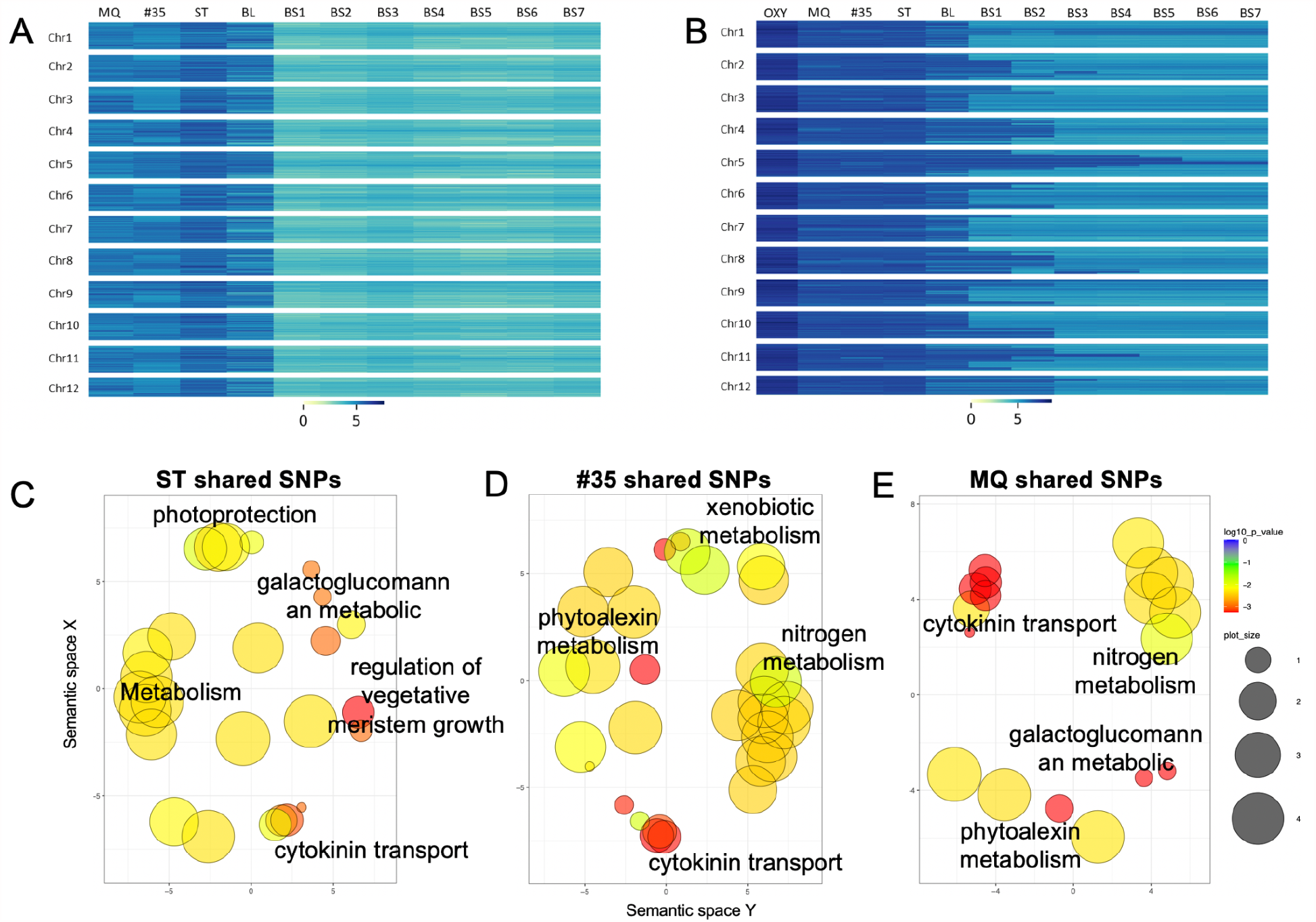
Regions of high and low SNP diversity in a wild selection and breeding-derived cranberry cultivars. A) Unique SNPs identified in early ‘bred’ cultivars of cranberry (#35, MQ, and ST) compared to the wild selection (BL) and a series of inbred lines (BS1-BS7) B) Shared SNPs identified in early cultivated lines of cranberry (#35, MQ, and ST) compared to Voxy, the wild selection (BL), and a series of inbred lines (BS1-BS7). Overrepresented GO terms from shared SNPs across cranberry cultivars C) ST, D) #35 and D) MQ plotted in semantic space.

Voxy had the greatest SNP diversity, with over 99% of the 250 kb bins across the genome containing a significant number of unique heterozygous SNPs. In fact, only two bins were found to have a significantly lower number of unique heterozygous SNPs for Voxy. ST was the next genotype to exhibit a relatively high amount of unique SNPs, with nearly 45% of the bins containing a significantly large number of unique SNPs and only 7% of the bins being significantly low in unique SNPs. The other Vmac cultivars had a reduced number of unique SNPs. #35 had a high unique SNP count in 16% of the bins, and a low unique SNP count in 28% of the bins. MQ had a high unique SNP count in 11% of the bins and a low unique SNP count in 31% of bins. BL had the lowest number of unique SNPs outside of the inbred line (BL-S5) with only 4% of the bins containing a significantly high count of unique SNPs, and a low unique SNP count in 79% of the bins across the genome. As expected, the 5th generation self of Ben Lear (BL-S5) had a very low number of unique SNPs, with only a single bin having a significantly high number of unique SNPs and over 99% of the bins being significantly devoid of unique heterozygous SNPs (Figure 5A; Table S9).

These variable regions represent standing variation in wild populations as well as regions of early domestication in this young crop. We next assessed the functional significance of these highly variable regions by examining the GO terms of the underlying genes (Table S10-12). The significant GO terms based on the genes in the variable regions of ST were related to plant architecture, metabolism, and environmental response (Figure 5C; Table S10). In contrast, #35 and MQ both had significant GO terms related to terpene based phytoalexins (pest inhibitory compounds), and nitrogen metabolism (Figure 5D,E; Table S11,12). #35 did have significant GO terms in xenobiotic metabolism (metabolism of foreign chemicals) that was not found in either ST or MQ, which must have been lost in the breeding and selection process leading to MQ.

Underlying these regions of higher variation, are genes that are under selection, representing potential breeding targets. We looked at the selective pressure on the genes among the cultivars to identify possible targets of improvement. We only found 18 genes under positive selection (Ka/Ks >1) between BL and ST consistent with these lines being either a wild selection (BL) or a 1st-generation breeding selection (ST) (Table S13). In contrast, #35 and MQ had 785 and 786 genes under positive selection (Ka/Ks >2) (Table S13), although there were no significant (P<0.01) GO terms associated with these genes. We looked at specific genes in these lists for genes that have been the targets of selection. #35 and MQ share 34% (266) genes under positive selection with genes associated with plant architecture and photomorphogenesis: *EPIDERMAL PATTERNING FACTOR-like* (*EPFL*), *SHOOT GRAVITROPISM* (*SGR*), *STEROL METHYLTRANSFERASE* (*SMT1*), and *HEMERA* (*HMR*). Moreover, MQ had additional genes under positive selection in the photomorphogenesis, flowering and circadian pathway: *LIGHT-DEPENDENT SHORT HYPOCOTYLS* (*LSH*), *ENHANCER OF AG-4* (*HUA2*) and *LHY*. In addition to being under positive selection, *LHY* is also tandemly duplicated in Vmac (Figure 3B), suggesting it may play an important role in the domestication of cranberry consistent with the selection pressure on core circadian genes in other crops [39].

## Discussion

Here we describe an updated genome assembly for the highly inbred reference cranberry accession Ben Lear (Vmac BL-S5) and a draft assembly for *V. oxycoccos* (Voxy) that is currently being used in our cranberry improvement program. The chromosome resolved Vmac genome confirmed the two WGD in the cranberry lineage that along with the more recent TDs have acted to shape the domesticated cranberry genome. Moreover, comparison with Voxy and more advanced selections of Vmac, revealed that response to the environment and plant architecture are under selection. The Vmac reference genome and Voxy draft genome will greatly facilitate current efforts to generate improved cranberry selections.

While we were preparing this manuscript a chromosome-resolved genome for a different cranberry accession (Stevens; ST) and a fragmented Voxy genome assembly were published [20]. It is exciting to see genomic resources emerge for this iconic North American crop and surely having a high quality genome for a second accession will refine our knowledge of cranberry biology. We compared our reference Vmac (BL-S5) genome assembly to the ST assembly and found that the two two were highly collinear, but consistent with the lower contig N50 length of the ST assembly, there is substantial telomere and repeat sequence missing (Figure S10). This is evidenced by non-linear regions in the dotplot, higher rate of non-syntenic orthologs and missing regions around the putative centromeres. A more thorough analysis of these two genomes will be the focus of future work.

We reported previously that the sugars associated with the anthocyanins are distinct between Vmac and Voxy [18]. Vmac contains primarily galactosides and arabinosides of the aglycones cyanidin and peonidin, while Voxy contains mostly glucosides of the same aglycones. This difference is important as it may affect antioxidant bioavailability. The sugar moiety is attached to the anthocyanidin by specific UDP-glucose:flavonoid 3-O-glycosyltransferases (UF3GT). Within the *UF3GT* is the highly conserved plant secondary product *glycosyltransferase* (PSPG) box. Amino acids in the PSPG box are reported to determine sugar specificity [40]. Specifically, the last amino acid in the PSPG box is reported to be specific for the sugar substrate with histidine conferring specificity for galactose and glutamine conferring specificity for glucose [41]. We have identified two Anthocyanidin 3-O-glycosyltransferases that exist distinctly in Voxy and Vmac (Vmac_055574 and Voxy_017508). The variant in Vmac contains histidine in the active site (Chr11-43267493), consistent with the galactosides found in Vmac anthocyanins, while Voxy has the glutamine amino acid associated with glucose specificity (Chr185-86547). Interestingly F1 interspecific hybrids of Vmac x Voxy have intermediate anthocyanin glycoside profiles, while about half the backcross (to Vmac) exhibit relatively high anthocyanin glucosides [18]. We identified other anthocyanidin 3-O-glycosyltransferases within the genomes of both Vmac and Voxy that may confer glycosylation of other flavonoids, e.g. flavonols conjugated to galactosides and arabinosides. However, there is only one location in the Voxy genome that contains the active site (which encodes the glutamine noted above), and only two in Vmac. Although 2 active sites are identified in Vmac, only one (that encodes the histidine) active site is located within a gene. Interestingly, just upstream of the annotated gene in Vmac ‘active’ gene there is an additional anthocyanidin 3-O-galactosyltransferase that is fragmented and lacks the complete active site, possibly explaining the dramatic differences in the anthocyanins between the two species.

Several genes that may play key roles in pathogen resistance have been identified being under selection pressure in the Vmac genomes. Both #35 and MQ show significant selection pressure for *PGIP2*, believed to play an important role in resistance to microbial colonization [42]. *SMT1*, a methyltransferase involved in sterol biosynthesis, is influential for innate immunity and the formation of *FLS2* receptor kinase clustering (flagellin sensing 2) [43]. *HIR3* is part of the hypersensitive response (HIR) gene family that has been shown to act in the defense of microbial infection as well as influencing cellular response during viral infection [44]. *LYK4* (Lysin motif domain receptor-like kinase 4) was shown to be an important plant defense component against fungal infection and is a key signalling component in plant chitin response [45]. Other genes found among the lines under selection pressure included *WRKY65, WRKY29, PALM1*, and *MLO*. While these specific genes were found in both #35 and MQ, there are several unique domains found in the wild relative Voxy that might offer further opportunity for incorporation into agricultural varieties.

Interestingly we also found defense related genes under selection pressure to be differentially expressed in the transcriptomes of other *Vaccinium* species during herbivory stress [46,47]. These genes included pleiotropic drug resistance transporter *ABCG36*, which provides pathogen resistance in Arabidopsis [48,49] and *FAH1*, also identified to be an important component of stress response in Arabidopsis [50]. Additionally, the serine/threonine-protein kinase *D6PKL2*, part of the auxin response pathway, is upregulated during herbivory in chickpea as well as bilberry [47,51]. In addition to pathogen resistance, several genes in the various Vmac lines were identified under selection pressure for stress tolerance. One of the key stressors includes drought stress, which is of particular importance to cultivated cranberry as a large portion of time in dormancy is spent under drought conditions. These genes include *CIPK2* [52], *AVP1* [53], *GAI* [54], *CPK20* [55], and *ABI4* [56]. Wax production on the fruit surface is an important trait for the protection against pathogens, UV damage, and for limiting moisture loss. We identified three genes that are related to wax production and UV protection that were under selection pressure in the Vmac lines. These genes included *PALM1* [57], *KCS2* [58], both relating to epicuticular wax production, and *MSH2* which is required for mitigation of UV-B light damage [59].

Changes in circadian components could be pressured due to the latitudes at which Voxy and Vmac reside. Ranges at higher latitudes could necessitate a greater amount of flexibility in the core circadian oscillator to compensate for large swings in light dark cycles throughout the course of the year. *LHY* is a key component of the core circadian oscillator, and *LHY* mutants have shown to have short photoperiods [60], while *PRR9* is an important component in the entrainment of the core oscillator to changes in photoperiod [61]. Taken together, changes in these circadian components could have larger downstream effects on seasonal flowering and fruit development [61–63].

Several other crop species have shown that circadian control and adaptation of photoperiod is important for both domestication and augmentation of desired traits, including flowering time, yield, and nutrient content [64–66]. The domestication of pea has been linked with variation in circadian genes for photoperiod response, including *HR* and *ELF3* [67], which are important interaction partners of *LHY* and *PRR9* for the regulation of photoperiod response [68,69]. This response allowed peas (*Pisum sativum*) to be cultivated at different latitudes, much like the differences between wild Voxy and Vmac. Additionally, flowering time expression is altered in domesticated cucumber when grown at varying latitudes [70]. Although we did not find the gene *FT*, a major influencer of flowering time, to be different between Vmac and Voxy, *LHY* is a key component in its regulation [62].

## Methods

### Plant growth

The cranberry cultivar Ben Lear (Vmac) was selected from the wild in Berlin, Wisconsin (43.9680° N, 88.9434° W) in 1901 [10]. To reduce heterozygosity, a fifth-generation selfing cycle inbred clone (F *≥* 0.97) of ‘Ben Lear’ designated BL-S5 (accession CNJ95-125-1) was selected for genome sequencing. The Voxy sequenced and used for hybridization with Vmac was collected near Gakona, Alaska (62.3019° N, 145.3019° W) in 1996 and designated NJ96-20 [15]. The hybrid (Vmac X Voxy) was the result of a cross (‘Stevens’ x NJ96-20) made by N. Vorsa in 1998, designated CNJ98-325-33. The ploidy of all cultivars and accessions used was confirmed by flow cytometry [22]. All plants were maintained in 6 inch pots containing sandy soil and fertilized with azalea mix for acidic plants. While maintained in a greenhouse, plants were allowed to winter chill and developed as ambient temperature increased.

### DNA extraction

Fresh leaf tissue of Vmac (CNJ95-125-1; BL-S5), Voxy (NJ96-20), and the hybrid (Vmac X Voxy, CNJ98-325-33) was stored in the dark for 3 days to reduce the polysaccharides. Tissue was then flash frozen in liquid nitrogen and ground into fine powder using mortar and pestle. High molecular weight (HMW) DNA was extracted with a modified CTAB protocol, optimized for cranberry [71]. HMW DNA was checked for quality on a Bioanalyzer (Agilent, Santa Clara, CA, USA) and length on a standard agarose gel. HMW DNA was used for library construction and sequencing on the long read Oxford Nanopore Technologies (ONT, Oxford, UK) platform and the Illumina (San Diego, CA) short read platform.

### Sequencing

HMW DNA was first sequenced on an ONT MinION sequencer to confirm quality for long read Nanopore sequencing. Unsheared HMW DNA was used to make ONT ligation-based libraries. Libraries were prepared starting with 1.5ug of DNA and following all other steps in ONT’s SQK-LSK109 protocol. Final libraries were loaded on an ONT flowcell (v9.4.1) and run on the GridION. Bases were called in real-time on the GridION using the flip-flop version of Guppy (v3.1). The resulting fastq files were concatenated (fail and pass) and used for downstream genome assembly steps. Illumina 2×150 bp paired end reads were generated for genome size estimates and polishing genome long read assemblies. Libraries for Illumina sequencing were prepared from HMW DNA using NEBnext (NEB, Beverly, MA) and sequenced on the Illumina NovaSeq (San Diego, CA). Illumina short reads for *V. macrocarpon* (CNJ95-125-1; BL-S5) were accessed from NCBI (PRJNA245813).

### Genome size prediction by k-mer frequency

Raw Illumina reads for *Vmac* (CNJ95-125-1; BL-S5; PRJNA245813), Voxy (NJ96-20) and the hybrid (Vmac X Voxy) were analyzed for k-mer frequency (k=31) using Jellyfish (count -C -s 8G -t 4 -m 31 and histo) [72]. Genome size was estimated and visualized using in house analysis scripts as well as GenomeScope [21]. While Vmac and Voxy had single peaks consistent with homozygous genomes, the hybrid had two peaks with the left peak bigger than the right peak, consistent with tetraploidy or the fact that the two genomes are distinct (Table S1; Figure S2).

### Genome assembly

Resulting ONT fastq files passing QC (fastq_pass) were assembled using our previously described long read assembly pipeline [24]. Briefly, fastq files were filtered by length for the longest 30x using a Illumina kmer-based genome size estimate [73]. The 30x fastq files were overlapped using minimap2 [74], the initial assembly was generated with miniasm [75], the resulting graph (gfa) was visually checked with Bandage [76], the assembly fasta was extracted from the gfa (awk ‘/^S/{print “>“$2”\n”$3}’ assembly_graph.gfa | fold >assembly_graph.fasta), the consensus was generated with three (3) iterative cycles of mapping the 30x reads back to the assembly with minimap2 followed by racon [77], and the final assembly was polished iteratively three times (3) using 2×150 bp paired-end Illumina reads mapped using minimap2 (>98% mapping) followed by pilon [78]. The resulting assemblies were assessed for traditional genome statistics including assessing genome completeness with Benchmarking Universal Single-Copy Orthologs (BUSCO) (Table 1; Table S2) [79]. The genome graphs were visualized using bandage (Figure 1) [76].

### Genome scaffolding

Cranberry (Vmac) is closely related (i.e. it is in the same genus) to *V. corymbosum* (highbush blueberry), which recently had an updated chromosome-scale genome release [25]. We leveraged the haplotype-resolved blueberry genome to assess the quality of our *V. macrocarpon* assembly by aligning our version 1 contig assembly (Vmac_v1) to haplotype 1 of blueberry at both the DNA level and the protein level. Vmac_v1 was aligned to Vcor_hap1 using minimap2 [74], and visualized the dotplot. Vmac_v1 was also aligned to *V. corymbosum* at the protein level using both CoGe [80], as well as MCscan (https://github.com/tanghaibao/jcvi/wiki/MCscan-(Python-version)) (Figure S3). Since the contig contiguity (N50 length) was 15 Mb for the Vmac_v1 assembly, which represents chromosome arms, we leveraged the synteny with the chromosome resolved Vcor_hap1 genome to orient Vmac_v1 contigs into super-scaffolds (chromosomes). The final Vmac_v2 assembly revealed several rearrangements between cranberry and blueberry, which were part of the original contig structure of Vmac_v1 (Figure S3). The Vmac_v2 chromosome assembly was versified using a high-density genetic map [26]. Linkage group (LG) specific anchors (>100 bp sequence) were created from the previous genome assembly [19] and used to validate order and orientation of Vmac_2 scaffolded contigs.

### Gene prediction and annotation

Genomes were first masked for repeat sequence before predicting protein coding genes. Repeat sequence was identified using the Extensive *de-novo* TE Annotator (EDTA) pipeline [31] (Table S3). ONT derived cDNA reads were aligned to the reference using minimap2 and then assembled into transcript models using Stringtie. We additionally leveraged two Illumina paired end cDNA libraries from SRA (SRR9047913, SRR1282422) as part of gene predictions. The soft masked genome was then used to predict protein coding genes using the funnannotate pipeline (https://funannotate.readthedocs.io/) leveraging the long read based transcript models and the illumina short read cDNA as empirical training data (Table 1). The resulting gene predictions were annotated using the eggNOG mapper [81].

### Disease Resistance

The complete CDS regions of Voxy and Vmac were analyzed through PRGdb’s DRAGO 2 API [35] to identify disease resistance motifs and further predict disease resistance gene annotations.

### Pollen Staining

Pollen stainability, with 1% lactophenol cotton blue stain, was employed to assess gamete fertility in Vmac and Voxy and the hybrid F1 interspecific progeny. Pollen was dusted on a microscope slide in a drop of stain and cover slipped. Pollen tetrads were observed at 400x magnification as described [82]. Pollen was determined to be viable if stained. Tetrads (pollen in *Vaccinium* spp. is shed with the 4 products of a pollen mother cell, as a tetrahedron). Tetrads were scored for 5 possible tetrad classes; four, three, two, one, or zero stained (viable) pollen grains.

### Gene family analysis

Gene family analysis was performed across several closely related species as well as several more distantly related species using OrthoFinder with default settings [83]. *Arabidopsis thaliana* (Araport11), *Amborella trichopoda* (v1) and *Vitis vinifera* (grape; v2.1) were accessed on Phytozome (https://phytozome-next.jgi.doe.gov/). The highbush blueberry (*Vaccinium corymbosum*) genome was accessed from CoGe (id34364) [25]; the rhododendron (*Rhododendron williamsianum*) genome was accessed from CoGe (id51210) [27], the persimmon (*Diospyros oleifera*) genome was accessed from http://persimmon.kazusa.or.jp [28], the tea (*Camellia sinensis*) genome was accessed from http://tpia.teaplant.org) [29] and the kiwi (*Actinidia chinensis*) genome was accessed from ftp://bioinfo.bti.cornell.edu/pub/kiwifruit [30]. Colored blocks in the figure generated (Fig. 2B) symbolize chromosomes or scaffolds while the lines (grey) symbolize syntenic regions between genomes. The Upset plot was generated from the orthogroup overlap file. The phylogenetic tree was constructed from the species_tree output from Orthofinder [83].

### Whole genome duplication (WGD) analysis

The genomes described in the gene family analysis were used for WGD analysis. Genomes for *A. thaliana*, A. *trichopoda*, grape, blueberry, rhododendron, persimmon, tea, and kiwi were aligned at the protein level using lastal in the MCscan python framework to calculate Ks and identify percentage of syntenic blocks across the genome pairs (https://github.com/tanghaibao/jcvi/wiki/MCscan-(Python-version)). Similar calculations were performed with genomes in CoGe [80] and FracBias was leveraged to confirm or identify syntenic block numbers underlying WGD events [84]. Karyotype figures were generated using MCscan python.

### Syntenic analysis

Syntenic analysis was performed between Vmac and Voxy using SyMAP v5 [85]. Data from the genome assembly as well as annotations of both genomes were imputed into SyMAP, though contigs of size less than 100 kb were not analyzed, while the otherwise default parameters were used to calculate synteny (-min_dots=7 -minScore=30 -minIdentity=70 -tileSize=10 -qMask=lower -maxIntron=10000). The subsequent analysis of overlapping syntenic blocks was performed with python scripting, where concurrent genome blocks of Voxy that overlapped the same location of Vmac were identified and gene annotation information of Voxy was pulled for further review.

### KaKs pressure

KaKs differences between Vmac and Voxy were calculated using gKaKs [86]. Genes under selection pressure (dN/dS>1) were cataloged for further analysis. GO terms were associated with genes by cross referencing the annotated gene name and the available data in the Uniprot database. Those GO terms associated with genes under selection pressure were collected for comparison. In addition, we compared two cultivars that are considered early domesticated varieties; Stevens (ST), #35, and a 3rd, later-domesticated variety Mullica Queen (MQ), with the wild reference, Ben Lear (BL). We identified differential GO terms between wild and domesticated lines, as well as several genes under selection pressure.

### Resequencing data analysis

Multiple generations of the BL inbreeding lines, as well as parents from several other lines important to the cranberry breeding program, were sequenced on the Illumina NGS platform. These included ‘Stevens’, ‘#35’, ‘Mullica Queen’, ‘Ben Lear’, a 5th generation self of ‘Ben Lear’ (BL-S5), and a wild accession of Voxy. Pedigree information of the resequenced parental lines can be found in [26]. Paired-end Illumina reads were aligned to the newly constructed Vmac reference genome (Vmac-v2) using BWA-MEM [87]. Reads were sorted and duplicate reads were removed from alignment files using samtools sort and rmdup respectively. SNPs were identified using samtools mpileup and bcftools call [88].

Further analysis was performed to identify comparative regions of high and low SNP density between lines. A script was generated in Python where heterozygous SNPs of each line, that were unique in both SNP position and nucleotide change for a single individual, were placed into 250,000 bp bins along the genome. The variant data from the genomes of Voxy as well as the genomes of the Ben Lear inbred lines (BL-S1 to BL-S7) were not used to determine uniqueness of SNPs compared to the rest of the Vmac lines since Voxy as well as the inbred lines would show disproportionate amounts of unique and non-unique SNPs respectively. Significant variation of unique SNP density was calculated through bootstrapping using the average SNP data of four representative varieties (Stevens, #35, Mullica Queen, and Ben Lear). 1,000 iterations were performed, where the aforementioned pooled SNPs were randomly assigned a bin, with the 95th percentile of bin maximums constituting the bounds of high SNP density and conversely, the 5th percentile of bin minimums constituting the bounds for low SNP density.

## Author contributions

TPM, JP, and NV conceived the study. TPM, KC and BA sequenced the genomes. TPM, JK, and NH analyzed data. JP, JK, and TPM wrote the paper.

## Data availability

The genomes are available through CoGe under genome ID 60226 and 60227 for Vmac and Voxy respectively. In addition, both genome assemblies with annotation can be found at the Genome Database for Vaccinium (https://www.vaccinium.org); Vmac accession number GDV21001 and Voxy accession number GDV21002. Assemblies and relevant read data can be located in NCBI under BioProject PRJNA738865; Vmac accession number SAMN19762178.

## Acknowledgements

We thank Shane Poplawski and Rachael Gominsky for DNA extraction and sequencing support. This research was funded through the following agencies and groups: USDA-NIFA-AFRI Grant 2017-67013-26215, New Jersey Blueberry and Cranberry Research Council and The Cranberry Institute. This work was supported by the Tang Genomics Fund to T.P.M.

## Supplemental Tables

**Table S1. Cranberry genome size estimates by k-mer frequency**.

**Table S2. Cranberry genome assembly BUSCO scores**.

**Table S3. Cranberry repeat prediction**.

**Table S4. Details of genes under positive selection between Vmac and Voxy.**

**Table S5. Gene ontology (GO) terms for tandem duplicated (TD) genes**.

**Table S6. Gene ontology (GO) terms for tandem duplicated (TD) genes unique to Vmac and Voxy**.

**Table S7. Disease resistant genes predicted by DRAGO2 in syntenic blocks between Vmac and Voxy**.

**Table S8. Orthogroup (OG) overrepresented gene ontology (GO)**.

**Table S9. High and low SNP region gene numbers for cranberry cultivars and inbred series**.

**Table S10. Stevens (ST) all high SNP regions significant GO terms.**

**Table S11. #35 all high SNP regions significant GO terms**.

**Table S12. MQ all high SNP regions significant GO terms**.

**Table S13. Genes under positive selection between important cranberry breeding cultivars and the wild selection Ben Lear (BL)**.

## Supplemental Figures

**Figure S1.**
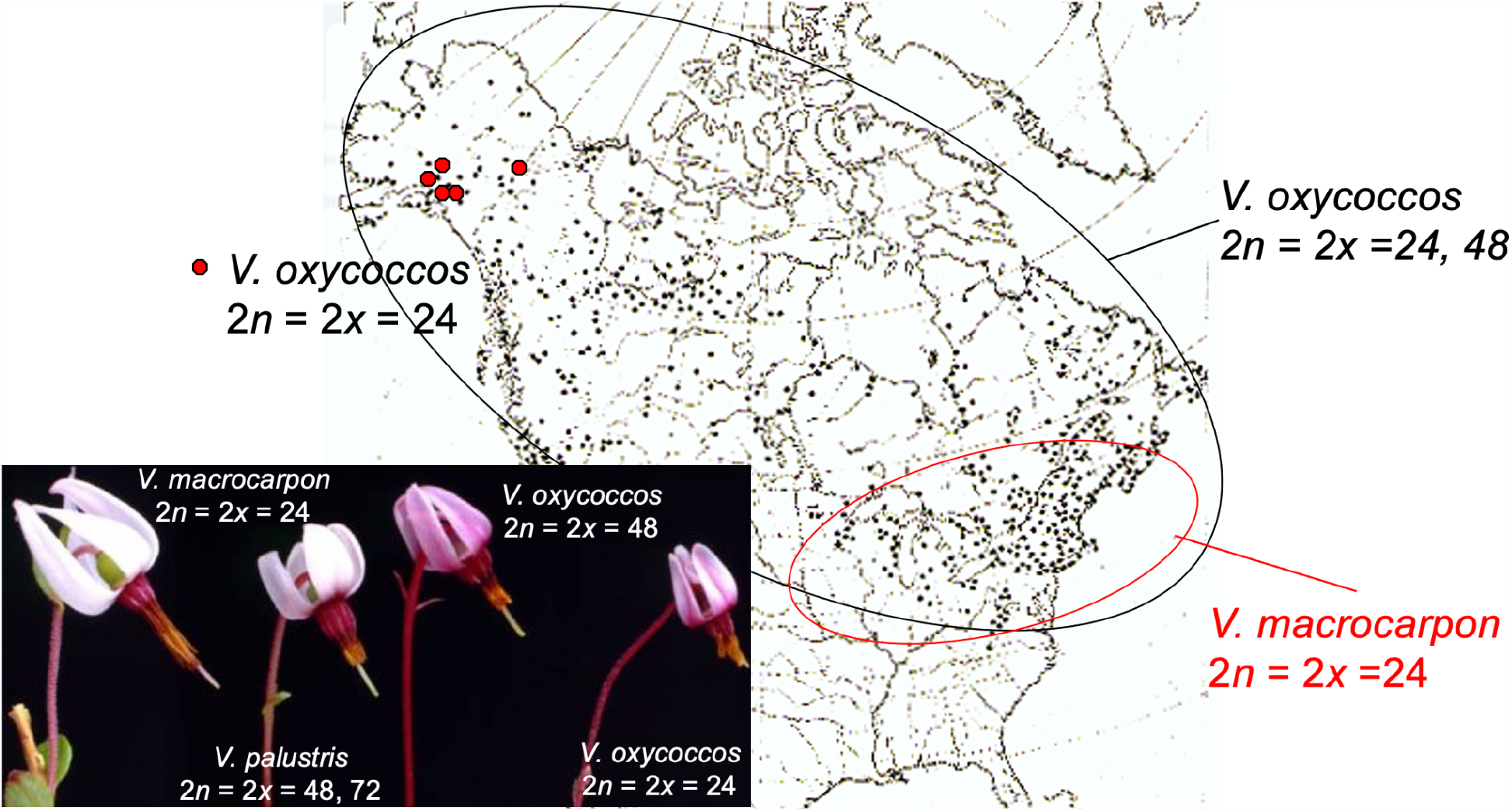
*V. macrocarpon* and *V. oxycoccos* distribution and flower size comparison. Diploid *V. macrocarpon* is found in the Northeastern parts of the United States (US), while the diploid *V. oxycoccos* is found in the Northwestern US and Canada.

**Figure S2.**
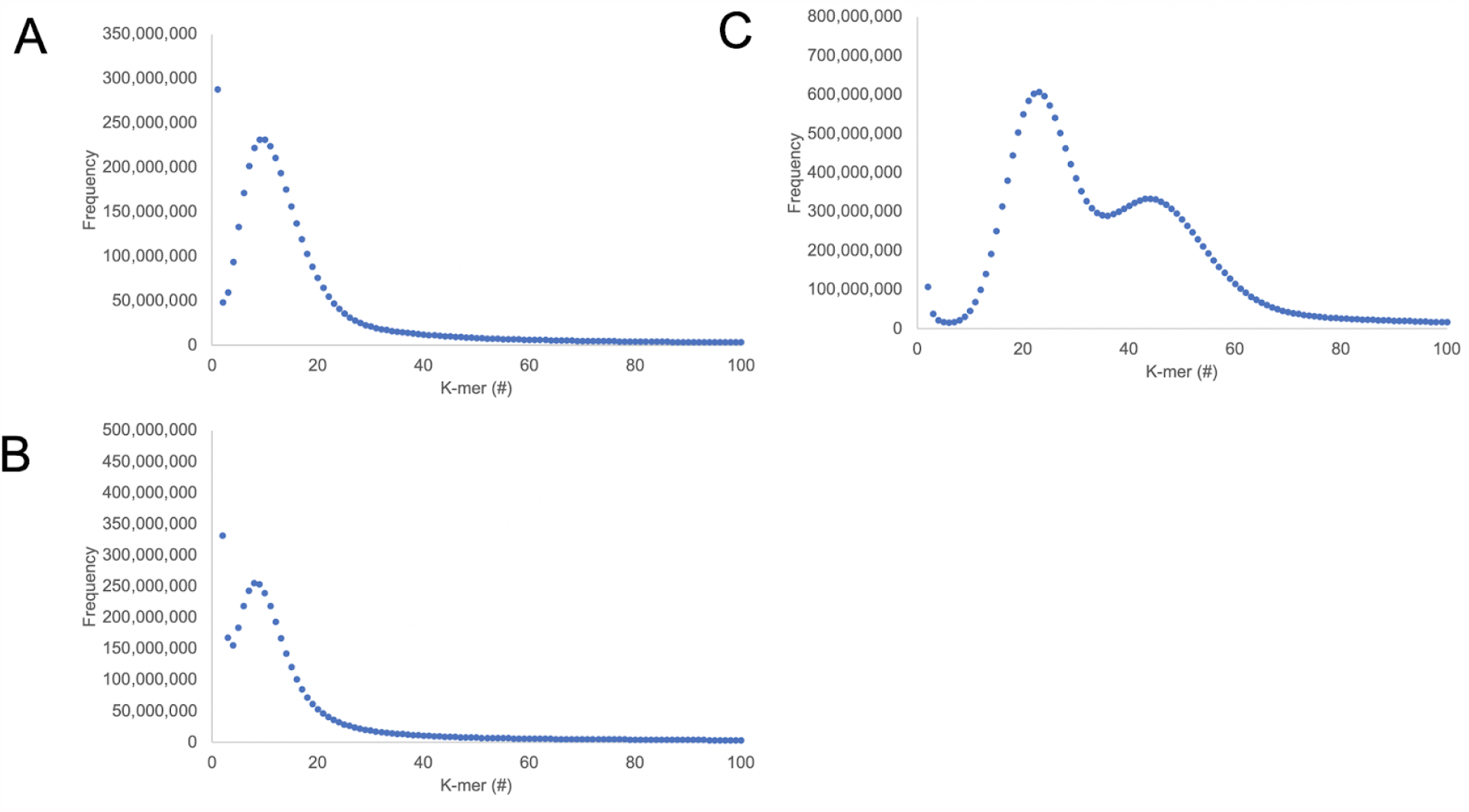
*V. macrocarpon* and *V. oxycoccos* genomes size estimated by K-mer. Genome sizes were estimated by K-mer (k=19) frequency using Illumina paired short reads (2×150 bp) for A) *V. macrocarpon* (Vmac), B) *V. oxycoccos* (Voxy), and C) the F1 hybrid. K-mers were counted with Jellyfish and histogram was plotted to find the peak.

**Figure S3.**
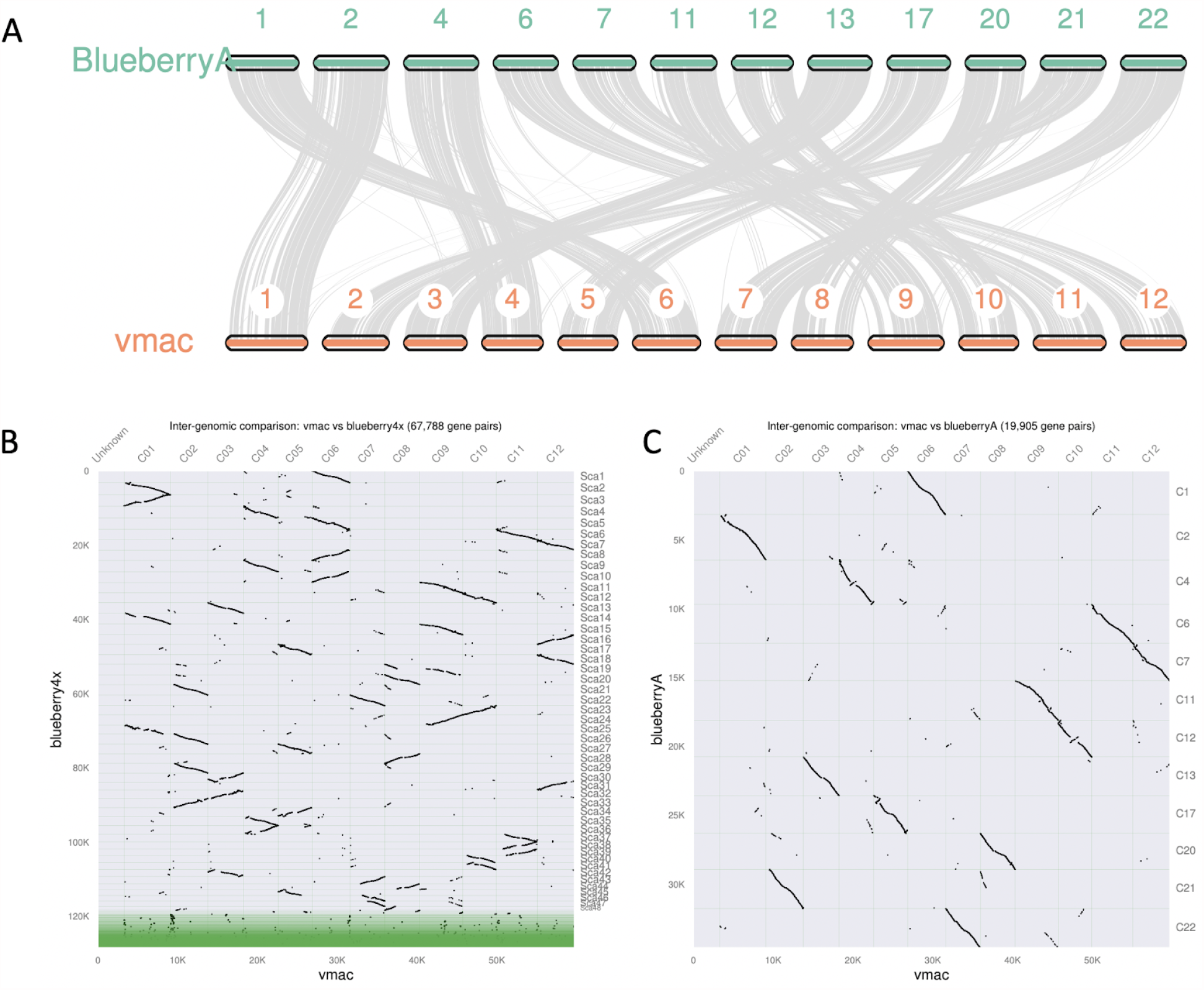
The *V. macrocarpon* (Vamc) genome is highly syntenic with chromosome-resolved blueberry (*V. corymbosum*) genome. A) Blueberry haplotype A (BlueberryA) was aligned to the Vmac assembly and are presented in the order of their assigned chromosome numbers. B) Dot plot based on protein alignments between the haplotype-resolved tetraploid blueberry (blueberry4x) and Vmac. C) Dot plot based on protein alignments between the blueberry haplotype A (blueberryA) and Vmac.

**Figure S4.**
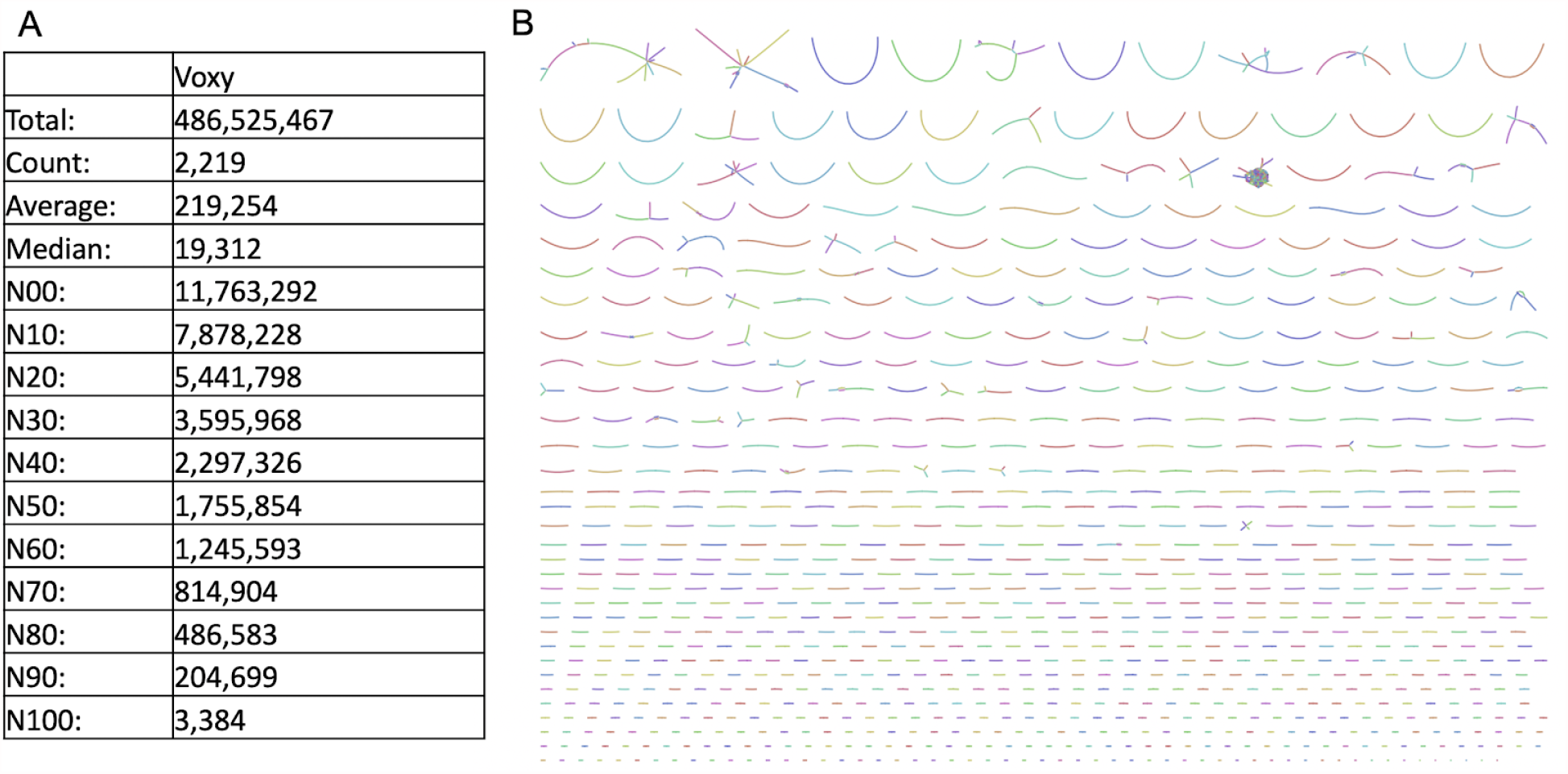
*V. oxycoccos* (Voxy) contig assembly and graph. A) Summary of the Voxy contig assembly statistics. The Voxy assembly was 486 Mb, had a N50 length of 1.8 Mb, with the longest contig being 11.8 Mb. B) The assembly graph of Voxy reveals low heterozygosity due to the lack of extensive branching.

**Figure S5.**
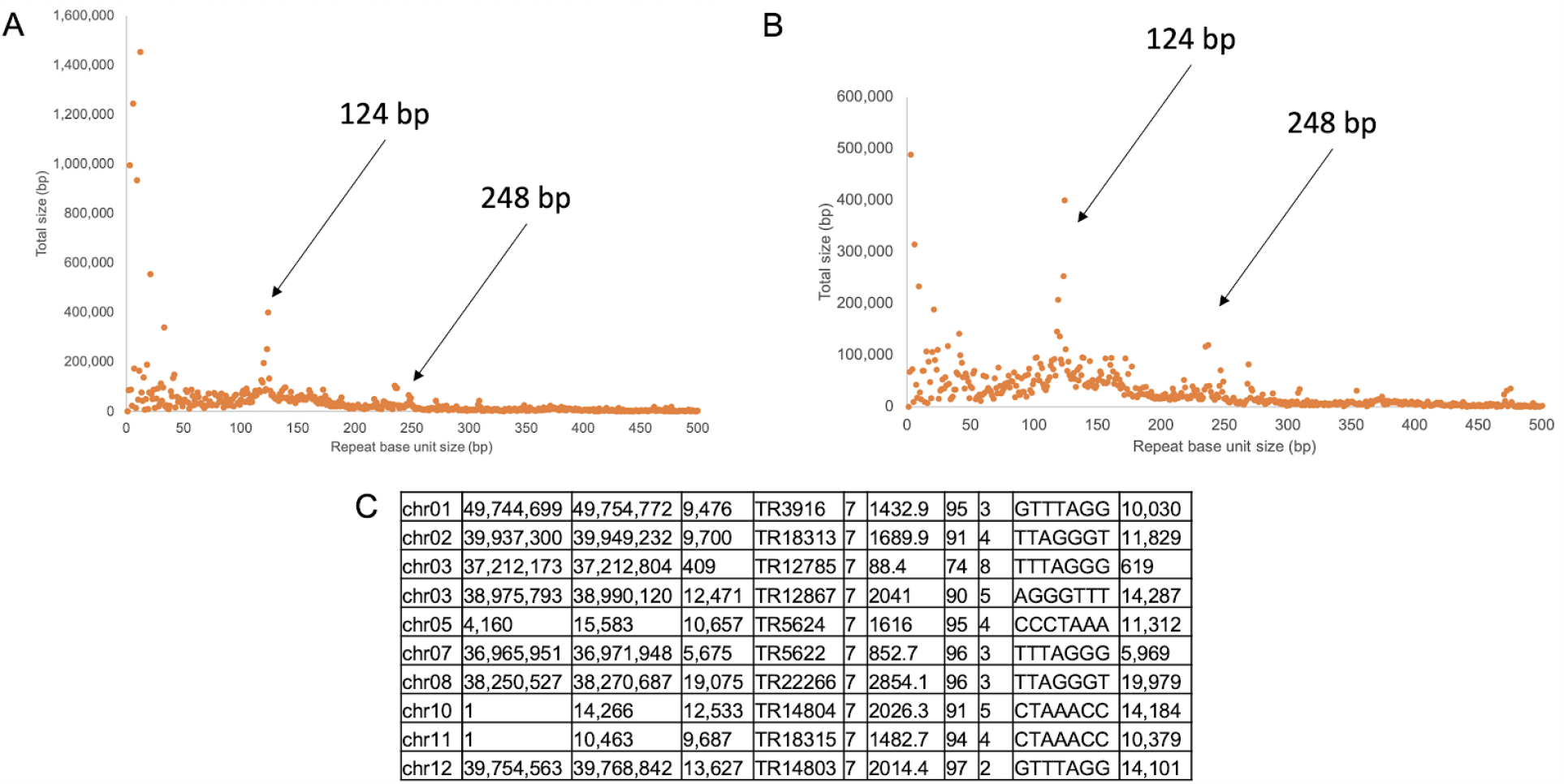
*V. macrocarpon* (Vmac) centromere and telomere arrays. A) Tandem repeats were identified using Tandem Repeat Finder (TRF) and plotted by repeat unit size, which revealed a 124 bp centromere base unit with a 248 bp higher repeat (HOR) consistent with a centromere array. B) A similar centromere array with a base unit of 124 bp and HOR of 248 bp was identified in Voxy. C) Telomere arrays with the 7 bp base unit (AAACCCT) were identified in the Vmac assembly, which revealed an average telomere length of 12 kb.

**Figure S6.**
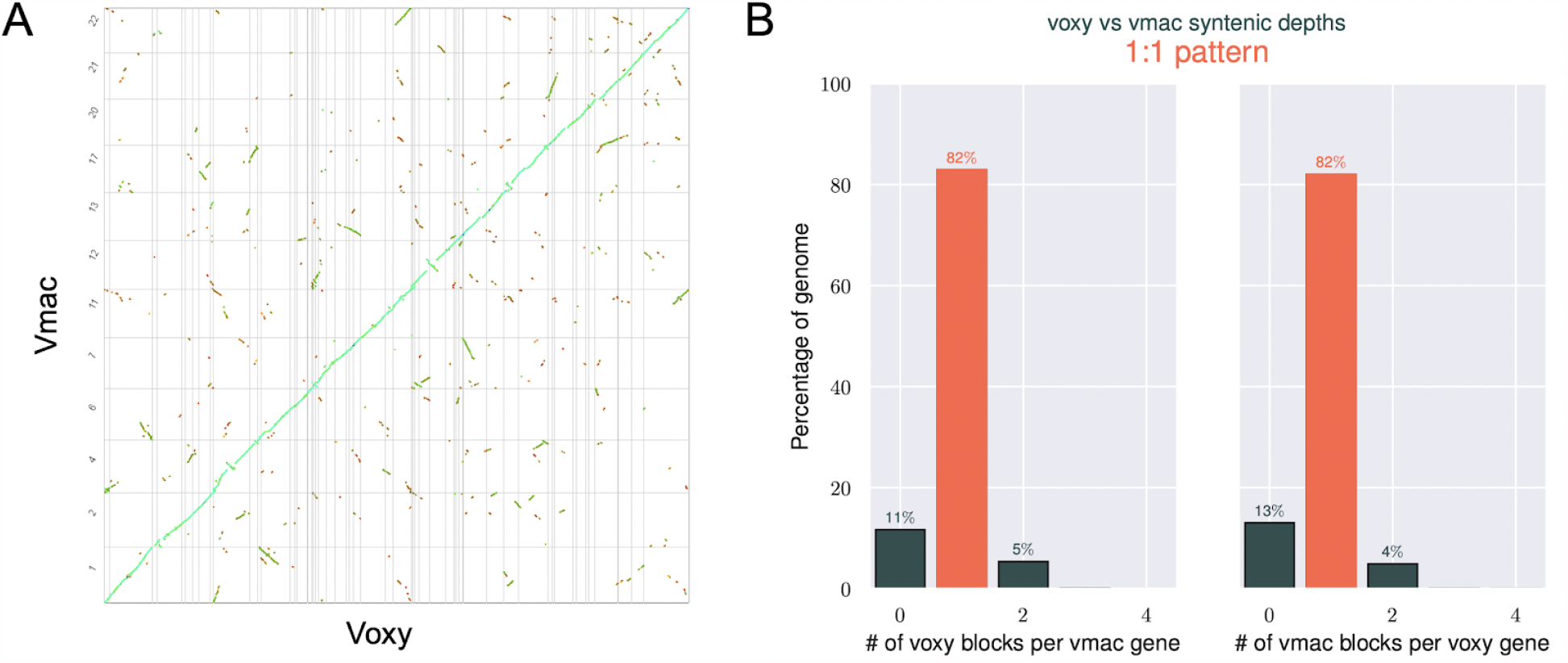
*V. macrocarpon* (Vmac) and *V. oxycoccos* (Voxy) are highly collinear. A) The Voxy scaffolds were aligned (protein) to the Vmac chromosomes revealing the two genomes are highly collinear with remnants of a recent whole genome duplication (WGD). Vertical and horizontal grey lines represent breaks in Chromosomes (Vmac) and scaffolds (Voxy) B) Syntenic depths between Vmac and Voxy suggest a 1:1 pattern, although there are remnants of a past WGD at 4-5%.

**Figure S7.**
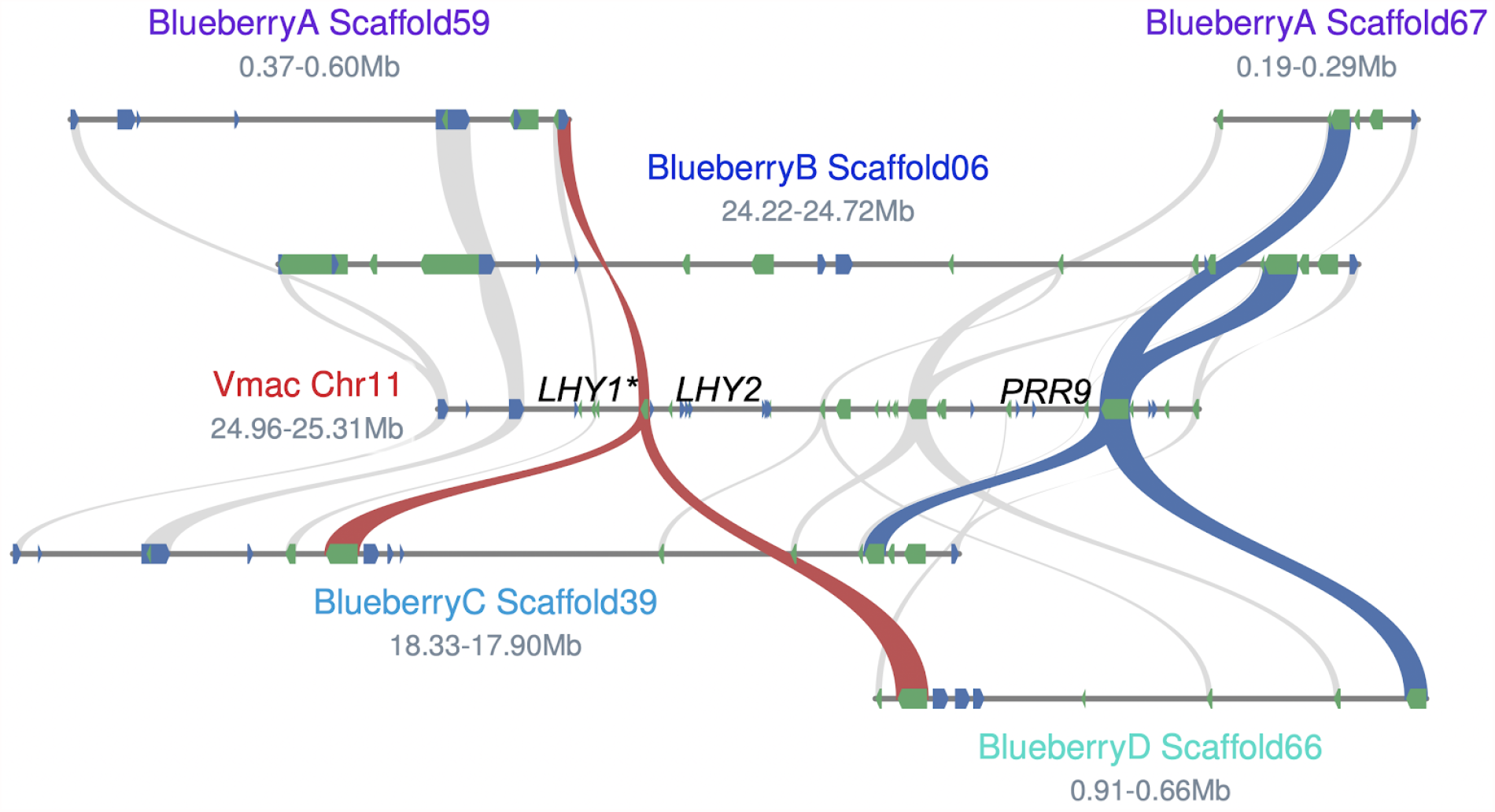
Tight linkage between core circadian clock genes is shared between cranberry and blueberry, but LHY tandem duplication (TD) is specific to cranberry. The haplotype-resolved blueberry genome was mapped to the Vmac genome to identify syntenic blocks (grey lines). Blueberry has the core circadian clock linkage of *LHY* (red lines)*-PRR9* (blue lines) on three of its haplotypes, but it has been lost on haplotype B on Scaffold6. The *LHY* tandem duplication is specific to the Vmac lineage since it is not found in Voxy (Figure 2) nor blueberry.

**Figure S8.**
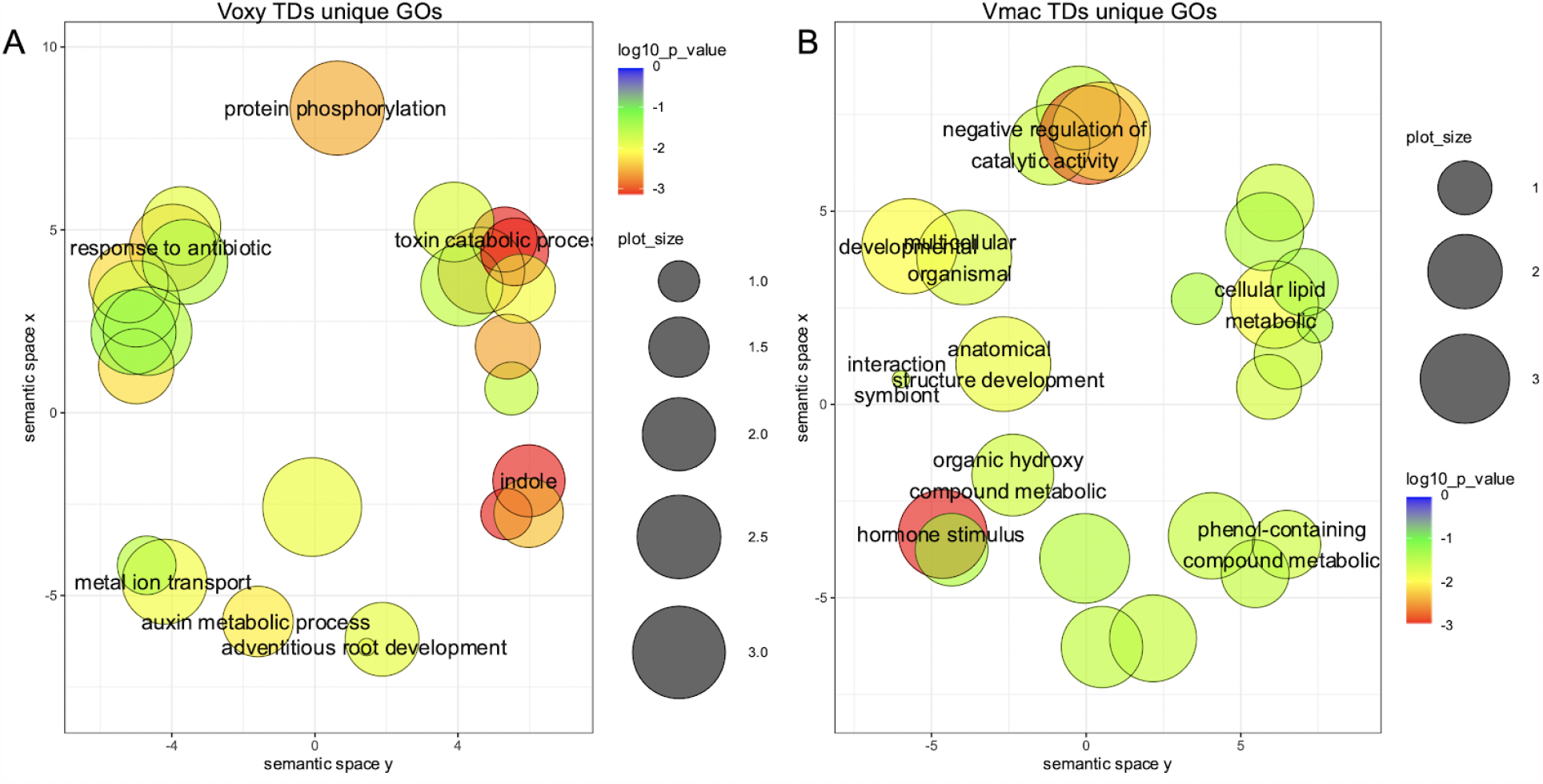
Overrepresented gene ontology (GO) terms found in unique tandem duplications (TDs) for *V. macrocarpon* (Vmac). A) *V. oxycoccos* (Voxy) TDs unique GOs are plotted in semantic space. B) Vmac TDs unique GOs are plotted in semantic space. Significance is colored with red being the most significant and blue the least significant. The size of the circle represents the number of elements.

**Figure S9.**
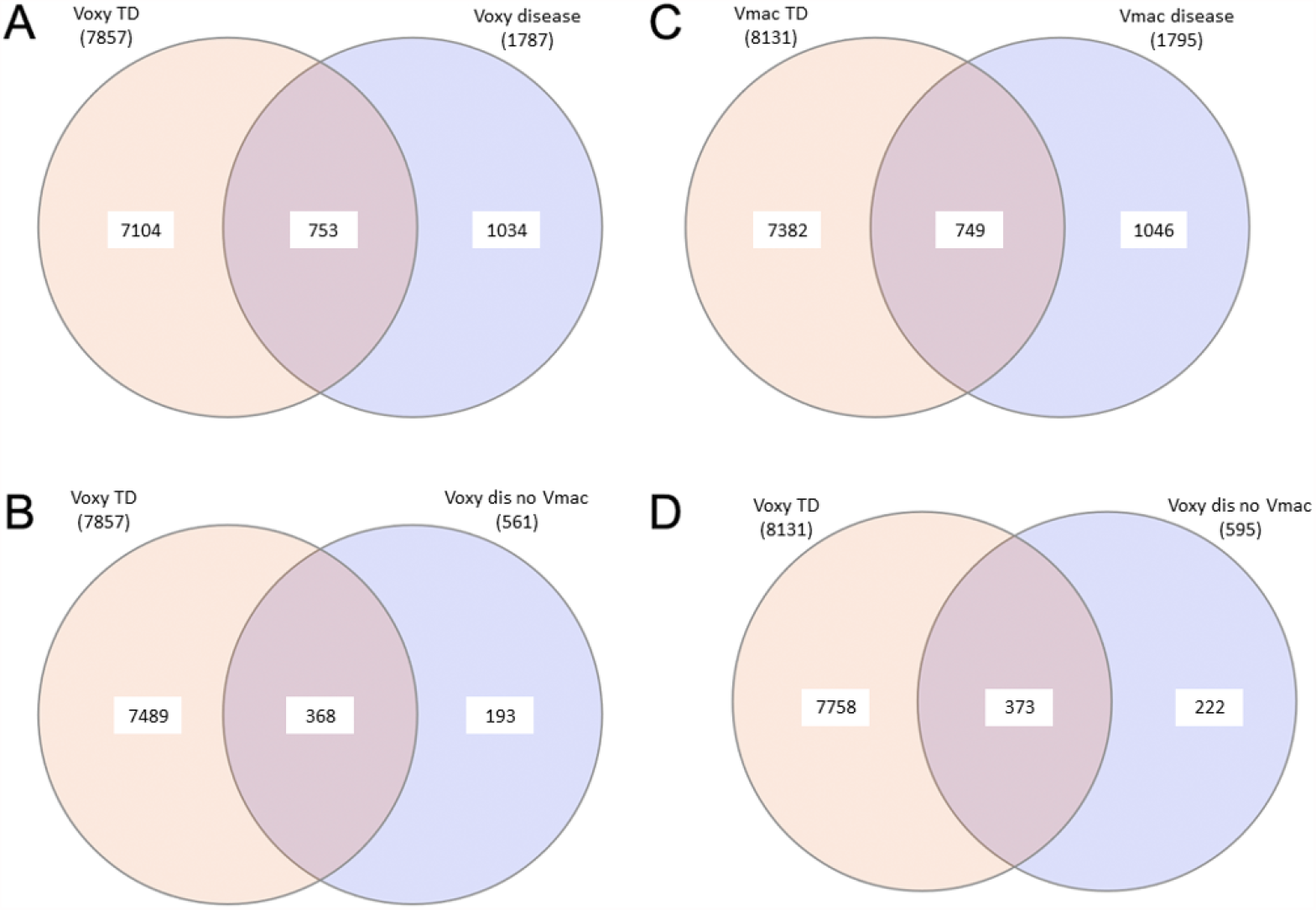
Venn diagrams of overlaps between predicted disease resistance genes and tandem duplications (TDs) in the *V. oxycoccos* (Voxy) and *V. macrocarpon* (Vmac) genomes. A) Voxy TD overlaps with predicted disease resistance genes, and B) disease resistance genes specific to Voxy (no syntenic ortholog in Vmac). C) Vmac TD overlaps with predicted disease resistance genes, and D) disease resistance genes specific to Vmac (no syntenic ortholog in Voxy).

**Figure S10.**
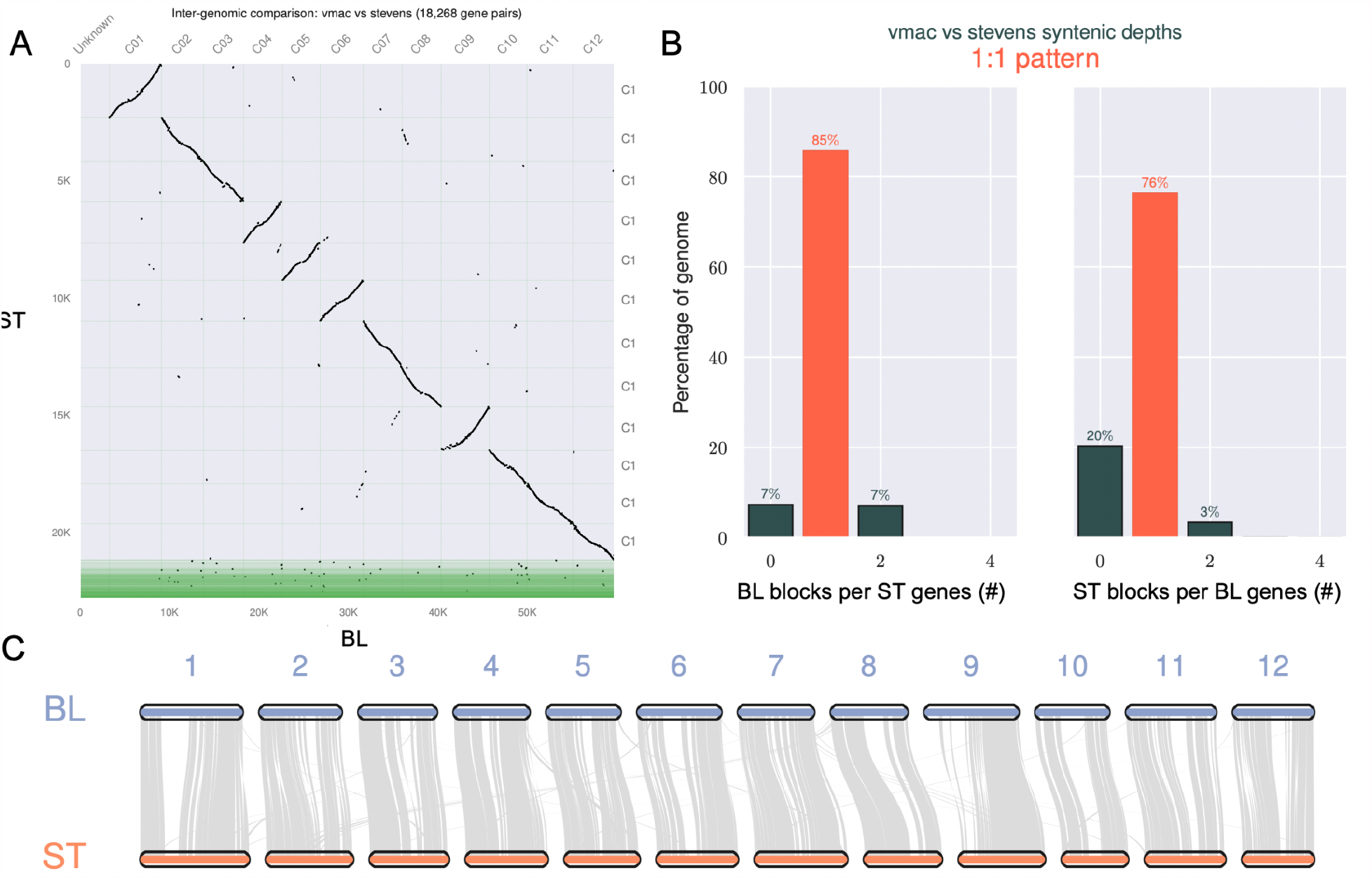
Comparison of the inbred *V. macrocarpon* Ben Lear S5 (BL) and the recently published Stevens (ST). A) Dotplot between BL and ST based on protein-protein comparisons reveals differences in chromosome size between the two access but high collinearity. Green area for ST are the contigs not included in the chromosomes. B) Syntenic ortholog patterns between BL and ST reveals that the ST genome is more fragmented than the BL genome due to more (20% vs 7%) genes with zero (0) syntenic blocks. C) Chromosome alignment between BL and ST with grey lines representing syntenic blocks. The missing regions between the two assemblies are centromere and repeat regions missing in ST.

